## Supplementary Note 1-8 for "OmicVerse: A single pipeline for exploring the entire transcriptome universe"

#### *OmicVerse is a unified resource for bulk-seq and single cell RNA-seq data analysis*

OmicVerse serves as a comprehensive resource for the analysis of both bulk-seq and single-cell RNA-seq data. The bulk RNA-seq analysis pipeline involves several essential steps. Initially, the data needs to be prepared in a raw matrix format (TPM or FPKM), which can be in txt, csv, tsv, or excel format. Subsequently, a meta matrix is required to label the sample features for subsequent analysis. OmicVerse offers a variety of data cleaning methods, including Deseq2 normalization[1], gene ID conversion, duplicate removal, and sample quality control[2]. Finally, OmicVerse facilitates downstream analysis such as differential expression analysis (pyDEG[1]), pathway enrichment analysis (pyGSEA[3]), weighted gene co-expression module construction (pyWGCNA[4]), protein interaction network analysis (pyPPI[5]), and bulk-seq to single-seq transformation (bulk2single[6]).

Similarly, the single-cell RNA-seq analysis pipeline consists of several steps. Initially, the data is organized within a data object using Scanpy[7] packages. OmicVerse provides a convenient function that automates normalization, scaling, log2 transformation, high variable gene filtering, dimensionality reduction, and differential expression analysis. Based on the results of pre-processed benchmark tests, omicverse provides a normalisation based on pearson residuals with highly variable gene extraction methods and optimises the logic of the scaled, progressive functions[8]. For more details please refer to our tutorial at https://omicverse.readthedocs.io/en/latest/Tutorials-single/t_preprocess/ . Additionally, OmicVerse currently implements models for various downstream analyses, including batch correction to integrate several samples (pyHarmony[9], pyCombat[10], scanorama[11]), automatic cell-type annotation (pySCSA[12]), trajectory inference (pyVIA[13], scLTNN[14]), pathway enrichment (pyGSEA[3], AUCell[15]), cell-cell interaction (CellPhoneDB[16]), factor analysis (pyMOFA[17]), single to spatial transformation (single2spatial[6]), metacell analysis (SEACells[18]) and drug response prediction (scDrug[19]).

Each model in OmicVerse is equipped with a simple and consistent application programming interface (API). These models rely on the pandas and AnnData formats to perform the analysis, allowing easy integration with Scanpy and scvi-tools [20]workflows. This seamless integration enables users to leverage the wider Python community for both single-cell RNA-seq and bulk RNA-seq analyses.

### Supplementary Note 2

#### *BulkTrajBlend reconstruction of mouse dentate gyrus neurons*

To assess the effectiveness of BulkTrajBlend in generating "interrupted" cells, we conducted an analysis utilizing single-cell RNA sequencing (scRNA-seq) data acquired from the dentate gyrus of the hippocampus in mice, in conjunction with bulk RNA-seq data. Within the single-cell data, we initially observed a neural differentiation trajectory spanning from neuronal intermediate progenitor cells (nIPC) to neuroblast to Granule immature to Granule mature. Interestingly, other cell types were seemingly disconnected from the nIPC trajectory, as illustrated in Extended Data Fig. 1g. A previous study had identified two distinct lineages, namely the nIPC to cortical projection neuron (CPN) and glial intermediate progenitor cell (gIPC) to oligodendrocyte progenitor cell (OPC) lineages[21]. Our primary objective was to validate the nIPC differentiation trajectory by linking OPC cells to nIPC cells within the "interruption" single-cell data.

Leveraging the capabilities of BulkTrajBlend, we initiated the deconvolution of the single-cell data, which consisted of 13 distinct cell types, using the bulk RNA-seq data from the dentate gyrus. We also filter out noisy subpopulations with unsupervised clustering. In the generated single cell profile, cell types that exhibited high expression patterns similar to those observed in the marker genes from the original single-cell profile, as evident in Extended Data Fig. 1a and Fig. 1b, respectively.

Subsequently, BulkTrajBlend quantified the overlap of cell types in the generated single-cell data using GNN-based Neural Overlapping Community Detection (NOCD), visually represented in the form of an adjacency matrix within a heatmap, as seen in Extended Data Fig. 1c and 1d. Within this generated single-cell data, we further characterized the overlapping cell community and specifically focused on the single-cell profile where OPCs were associated with nIPC in overlapping cell community, as depicted in Extended Data Fig. 1e-1f.

To assess false positives of predicted cells, we introduced the error overlap rate as a judgement of false positives of predicted cells, which is based on the following assumptions:

1, we assumed that a particular type of cell deconvolved from Bulk has a steady state, and its particular cell in a non-steady state is regarded as a transition cell

2, We assume that the transition cells have the characteristics of both types of cells and are seen as overlapping communities in terms of community, and that this fraction of transition cells can be captured using the GNN.

We define False Overlap Rate (FOR) as the judgement of false positive rate, defined as follows: if the overlapping community OC, in which a certain type of cell is located, is not contained in his unique original community UC, we denote it as False Overlap FO, and the formula of False Overlap Rate FOR is defined as follows:

$$FOR = \frac{Number_{FO}}{Number_{OC}}$$

We found that the FOR of nIPC, Cck-TOX, and Neuroblast is higher than 0.3, while the OPC we used for interpolation is 0.21, indicating that wrongly overlapping cells only account for 20% of the overlapping communities, which we used for back-interpolation (Extended Data Fig.1g).

We then seamlessly integrated this selected data into the original single-cell data from the dentate gyrus. Remarkably, our analysis revealed that the OPCs in the integrated scRNA-seq data of the dentate gyrus were positioned on the differentiated branch of nIPC, as illustrated in Extended Data Fig. 1h-1i. This intriguing observation demonstrated the efficacy of BulkTrajBlend in generating a continuous cell trajectory, successfully addressing the challenge of "interrupted" cells within the data.

This analysis not only underscores the power of BulkTrajBlend but also contributes to a deeper understanding of the cellular dynamics and differentiation processes in the dentate gyrus.

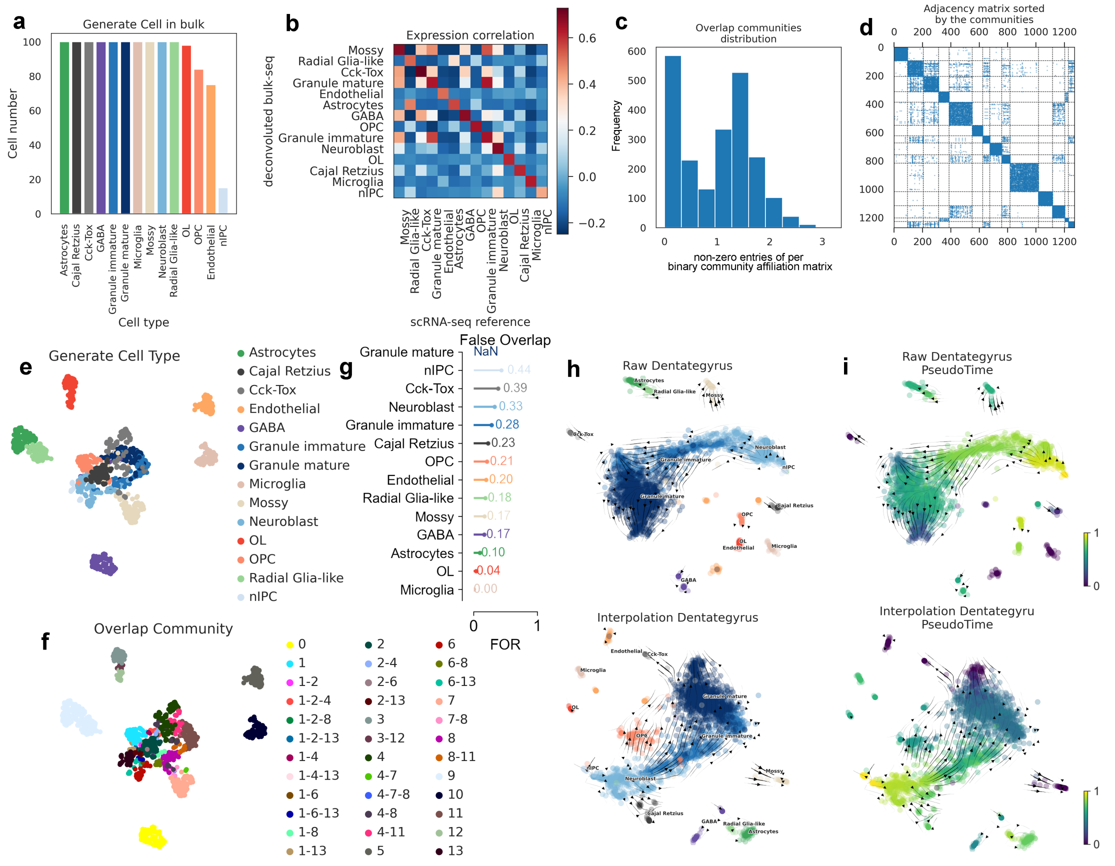

Extended Data Fig 1 | Application of BulkTrajBlend in Dentate Gyrus Neurogenesis.

(a) - Number of Cell Types in scRNA-seq Data Generated from Bulk RNA-seq using BulkTrajBlend.

(b) - Pairwise Expression Correlation of Cell-Type-Specific Marker Genes between Single Cells Generated by BulkTrajBlend and the Single-Cell Reference for Dentate Gyrus Neurogenesis.

(c) - Distribution of Overlapping Cell Communities in scRNA-seq Data Generated.

(d) - Heatmap displaying each cell's assignment to different cell communities in the adjacency matrix obtained from Graph Neural Networks (GNN).
(e) - UMAP Visualization of Cell Types in scRNA-seq Data Generated.
(f) - UMAP Visualization of Overlapping Cell Type Communities in scRNA-seq Data Generated.
(g) - False Overlap Rate in generated cell types, NaN represents the absence of major cell clusters of such cells in overlapping communities
(h) - Force-Directed Graph Comparison between Raw scRNA-seq (Upper) and Interpolated scRNA-seq (Bottom) Data for Dentate Gyrus Neurogenesis, Color-Coded by Cell Type.
(i) - Pseudo-time Analysis of Raw scRNA-seq (Upper) and Interpolated scRNA-seq (Bottom) Data for Dentate Gyrus Neurogenesis.

#### *Systematic benchmarking of interrupt performance*

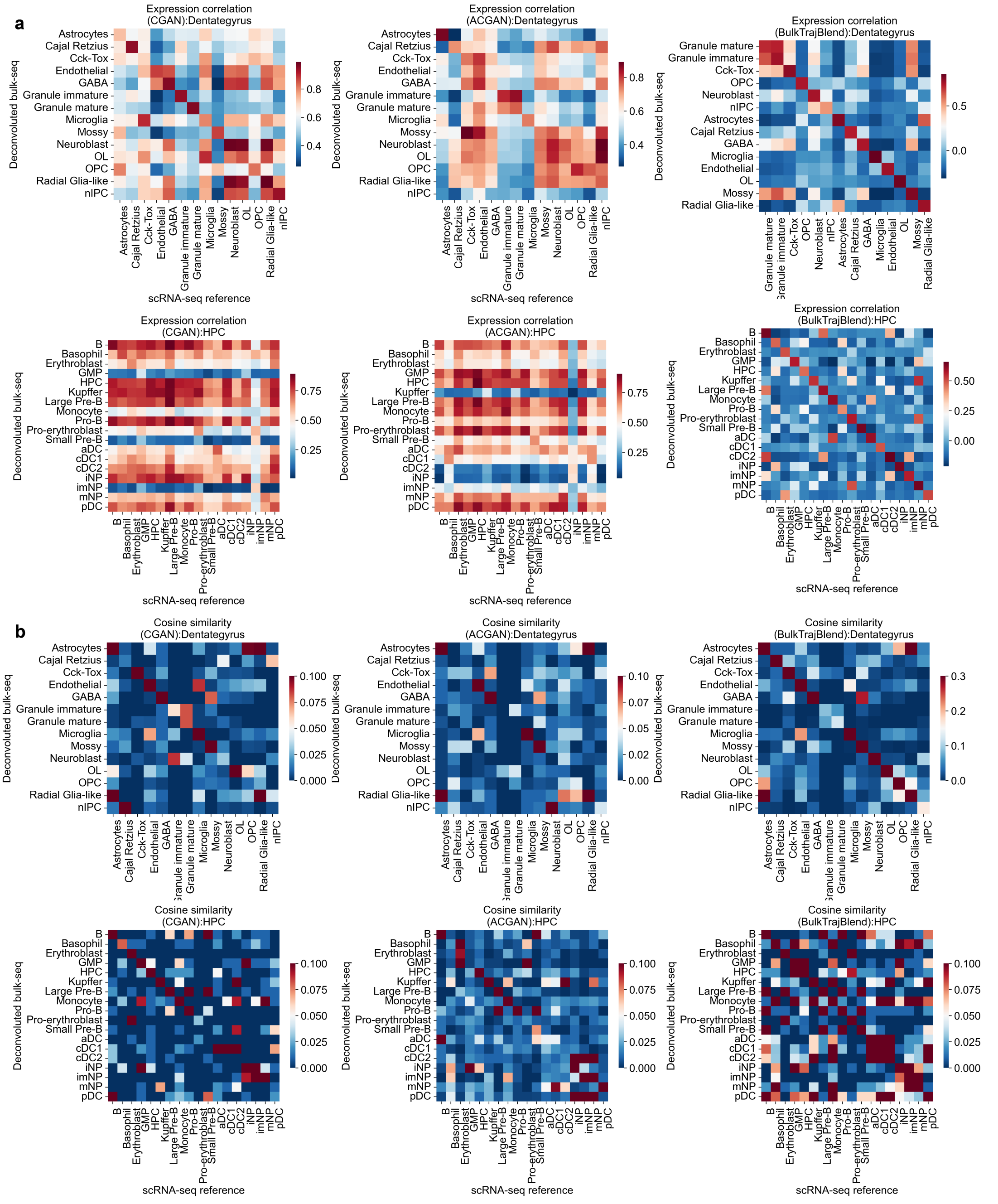

Extended Data Fig 2 | Assessment of Similarity in Generated Single-Cell Profiles and Raw Single-Cell Profiles

(a) - Correlation of Expression Trends of Marker Genes in Reference Single Cells between the Reference Single-Cell Profile and the Generated Single-Cell Profile. The upper panel represents the Dentate Gyrus profile, and the lower panel represents the Hematopoietic profile. From left to right, the methods examined are CGAN, ACGAN, and BulkTrajBlend.

(b) - Similarity Analysis of Marker Genes in the Reference Single-Cell Profile and Marker Genes in the Generated Single-Cell Profile. The upper panel represents the Dentate Gyrus profile, and the lower panel represents the Hematopoietic profile. From left to right, the methods assessed are CGAN,

ACGAN, and BulkTrajBlend.

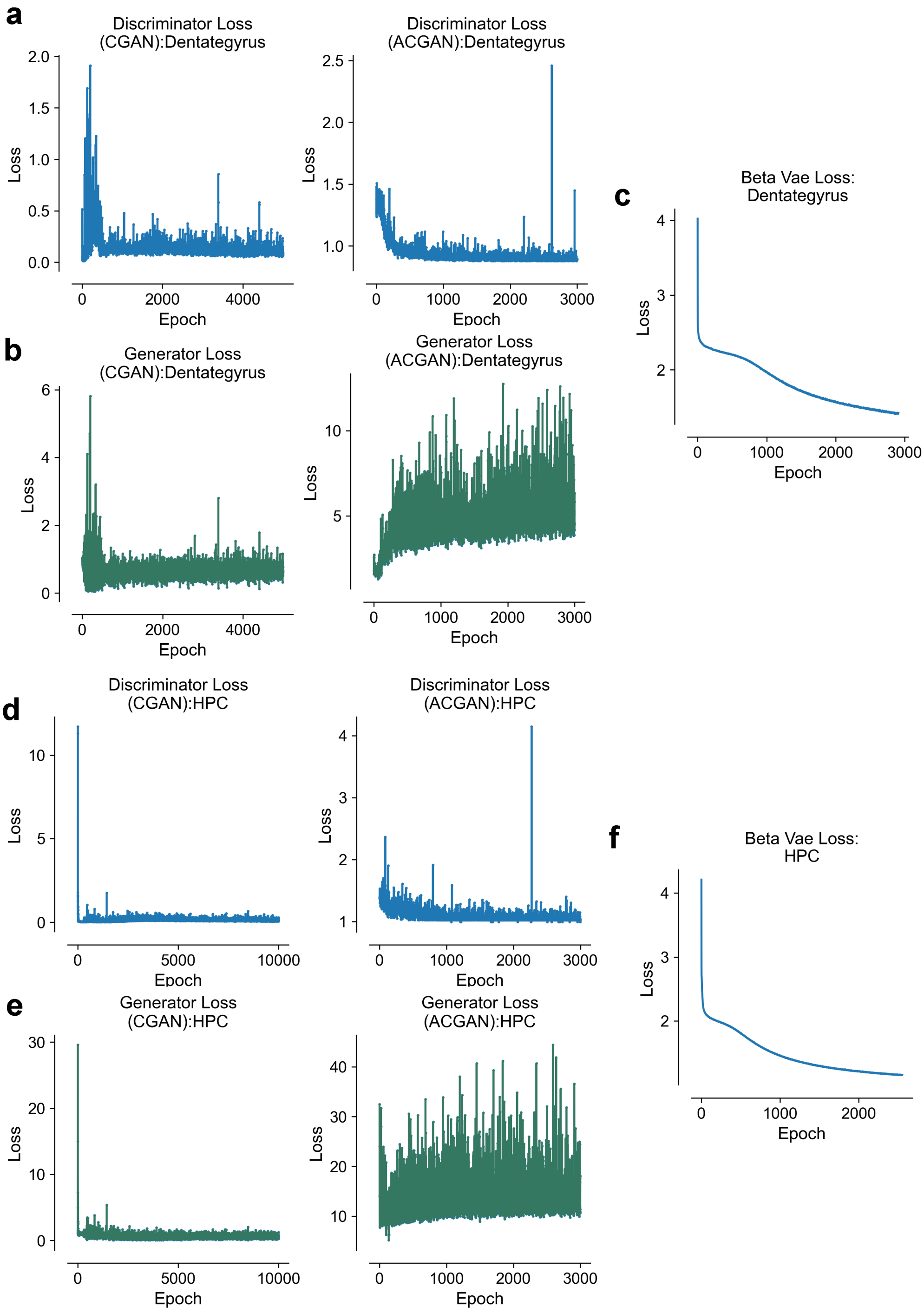

Extended Data Fig 3 | Training Process of Generation Models

(a) - Discriminator Loss on the Dentate Gyrus Dataset, where the horizontal axis represents training epochs, and the vertical axis represents training losses. The methods presented from left to right are CGAN and ACGAN.

(b) - Generator Losses on the Dentate Gyrus Dataset. The methods presented from left to right are CGAN and ACGAN.

(c) - Beta-VAE Loss on the Dentate Gyrus Dataset.

(d) - Discriminator Loss on the Hematopoietic Dataset, with the horizontal axis representing training epochs. The methods presented from left to right are CGAN and ACGAN.

(e) - Generator Losses on the Hematopoietic Dataset. The methods presented from left to right are CGAN and ACGAN.

(f) - Beta-VAE Loss on the Hematopoietic Dataset..

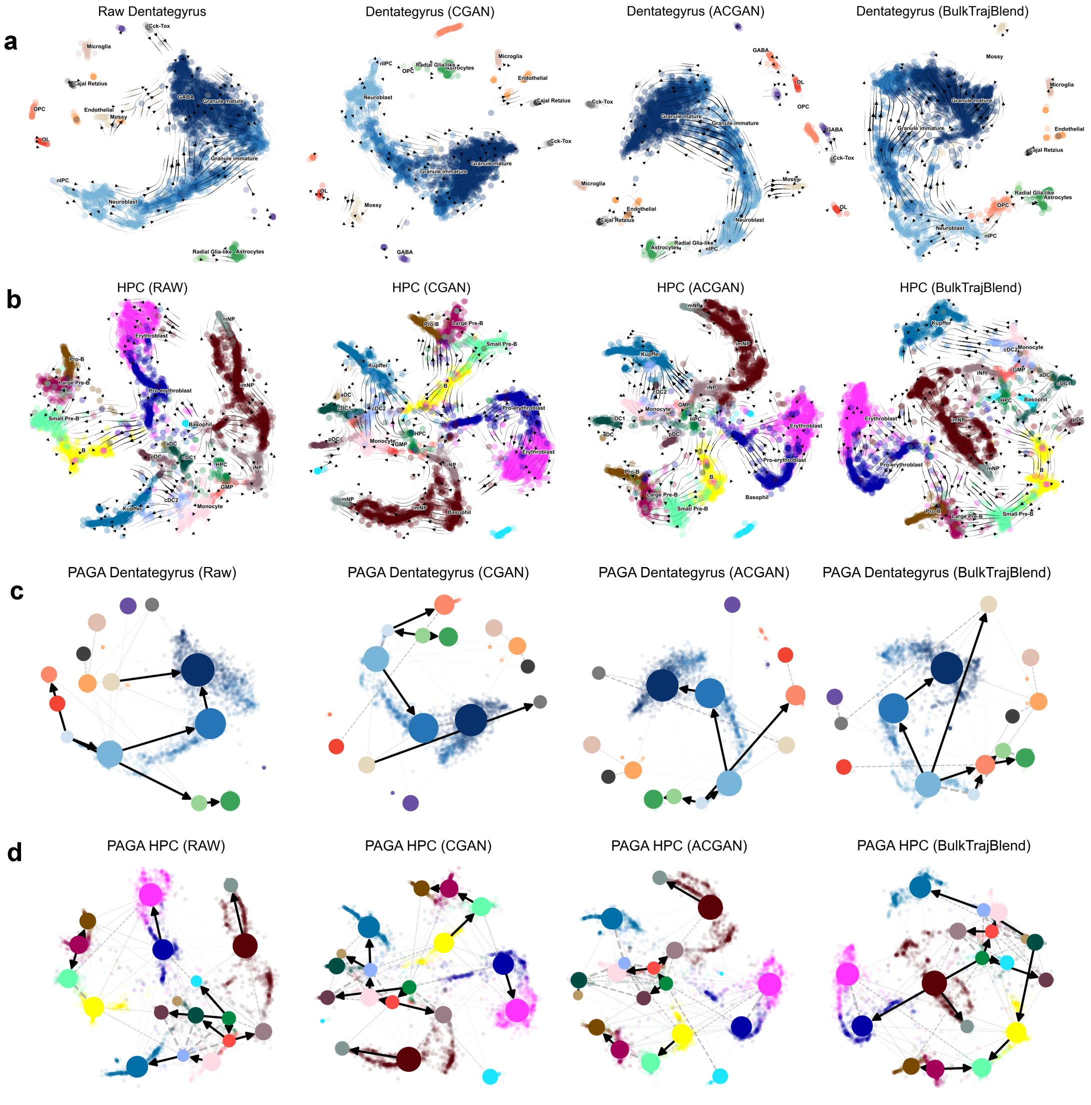

Extended Data Fig 4 | UMAP Visualization of Single-Cell RNA Sequencing (scRNA-seq) Data

(a) - UMAP plots illustrating the flow trend of cell developmental trajectories for neural progenitor cells (nIPC) within the Dentate Gyrus. Representations are provided for the RAW, CGAN, ACGAN, and BulkTrajBlend methods.

(b) - UMAP plots illustrating the flow trend of cell developmental trajectories for hematopoietic stem cells (HSC) within the Hematopoietic system. Data is presented for the RAW, CGAN, ACGAN, and BulkTrajBlend methods.

(c) - Directed Cell State Transfer Graph within the Trajectory of the PAGA Graph for the Dentate Gyrus. Results are displayed for the RAW, CGAN, ACGAN, and BulkTrajBlend approaches.

(d) - Directed Cell State Transfer Graph within the Trajectory of the PAGA Graph for the Hematopoietic System. Findings are showcased for the RAW, CGAN, ACGAN, and BulkTrajBlend methodologies.

### Supplementary Note 3

#### *BulkTrajBlend Efficiently Reconstructs Cell Developmental Trajectories in simulated single-cell profile*

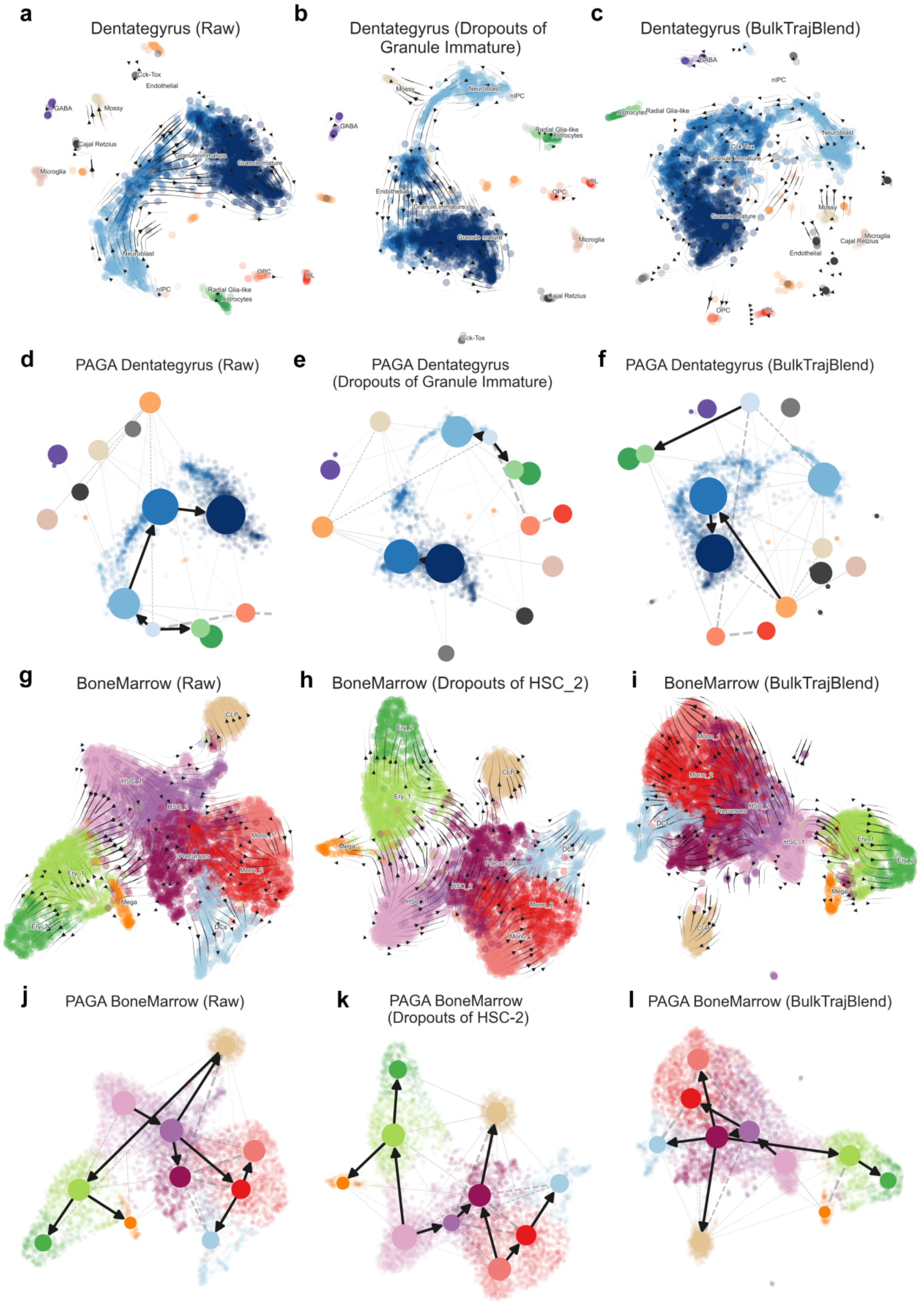

Extended Data Fig 5 | UMAP Visualization and Directed Graphs of Dentate Gyrus and Bone Marrow scRNA-seq Data

(a-c) - Velocity stream representations from left to right: (a) raw Dentate Gyrus dataset, (b) dataset with simulated cell dropouts, and (c) dataset interpolated with BulkTrajBlend to address dropouts estimated by pyVIA. The UMAP embedding is color-coded by cell type based on original cluster annotations.
(d-f) - Superimposed directed graphs on the UMAP embedding from left to right: (d) raw Dentate Gyrus dataset, (e) dataset with simulated cell dropouts, and (f) dataset interpolated with BulkTrajBlend to handle dropouts estimated by pyVIA.
(g-i) - Velocity stream visualizations from left to right: (g) raw Bone Marrow dataset, (h) dataset with simulated cell dropouts, and (i) dataset interpolated with BulkTrajBlend to mitigate dropouts estimated by pyVIA. The UMAP embedding is color-coded by cell type following the original cluster annotations.
(j-l) - Overlay of directed graphs onto the UMAP embedding from left to right: (j) raw Bone Marrow dataset, (k) dataset with simulated cell dropouts, and (l) dataset interpolated with BulkTrajBlend to address dropouts estimated by pyVIA.

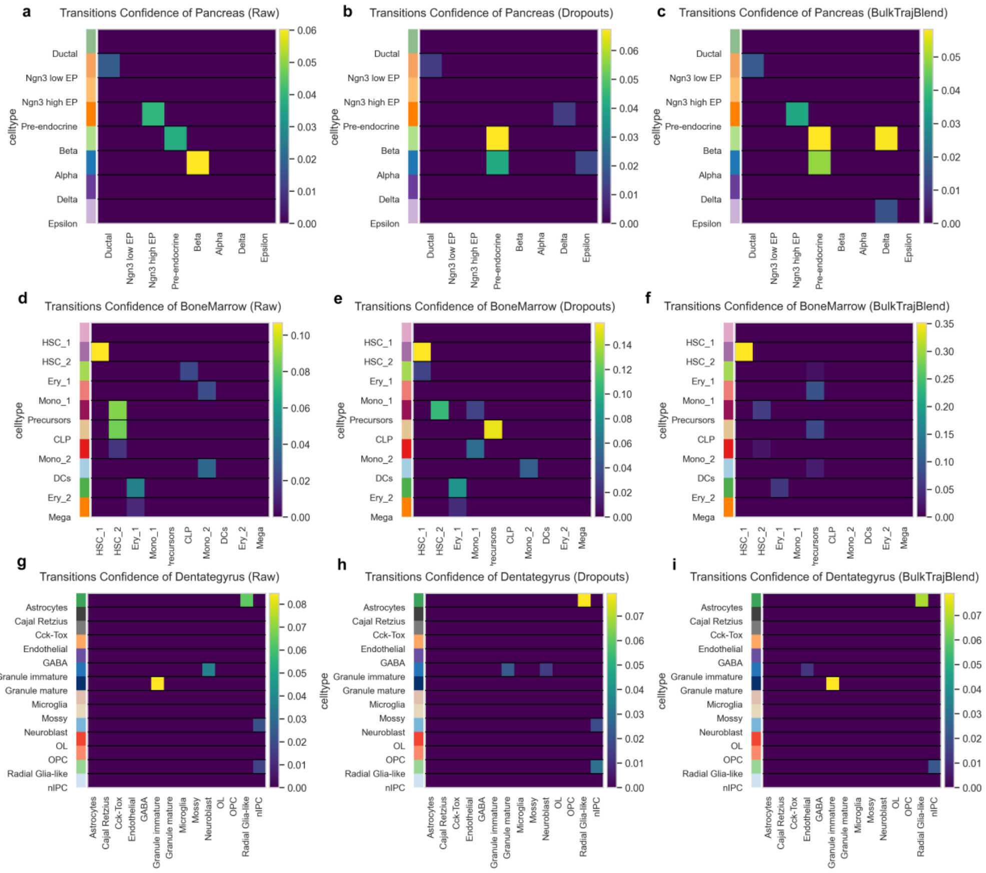

Extended Data Fig 6 | Analysis of Transition Confidence in scRNA-seq Data
(a-c) - Pancreas Dataset: Transition confidence analysis for the raw dataset, simulated cell dropouts, and BulkTrajBlend interpolation. The color scheme reflects the confidence values.
(d-f) - Bone Marrow Dataset: Transition confidence analysis for the raw dataset, simulated dropouts, and BulkTrajBlend interpolation.
(g-i) - Dentate Gyrus Dataset: Transition confidence analysis for the raw dataset, simulated cell dropouts, and BulkTrajBlend interpolation.

### Supplementary Note 4

#### *OmicVerse provides a comprehensive analysis platform for bulk RNA-seq data.*

Bulk RNA sequencing is widely used for transcriptomic analysis of pooled cell populations, tissue sections, or biopsies[2]. While it is commonly employed to measure gene expression patterns, isoform expression, alternative splicing, and single-nucleotide polymorphisms, RNA-seq contains additional valuable biological information. This includes details on copy number alterations, microbial contamination, transposable elements, cell type deconvolution, and the presence of neoantigens. Recent advancements in bioinformatic algorithms have made it possible to extract this information from bulk RNA-seq data, expanding its scope.

#### *Methods*

The dataset related to Alzheimer's disease, as previously described in a Nature Genetics publication, was obtained from the Gene Expression Omnibus (Accession ID: GSE174367). Associated metadata were also retrieved from the National Center for Biotechnology Information (NCBI) using the same Accession ID. To preprocess the bulk RNA-seq data, we performed two crucial steps: (1) the removal of duplicate gene IDs and (2) data normalization followed by logarithmic transformation. Within the metadata, we classified the samples into two groups: 'Normal - No Pathology Detected' as the control group and 'Alzheimer's disease' as the treatment group. This classification was stored in the `Neuropath.Dx.1` field of the metadata. The control group consisted of 8 samples, while the treatment group comprised 44 samples.

To visualize the similarity between samples, we employed Principal Component Analysis (PCA). This analysis allowed us to calculate Pearson correlation coefficients.

In the pyDEG analysis, we specified the following parameters: a fold change threshold of 0.15, a p-value threshold of 0.05, and the utilization of DESeq2 normalization to mitigate batch effects. The ` omicverse.bulk.pyDEG.plot_volcano` and ` omicverse.bulk.pyDEG.plot_boxplot` functions were used to visualize the differentially expressed genes. Specifically, the top 10 genes with the highest fold changes were selected and displayed in the boxplot.

For the pyGSEA analysis, we employed all 16,504 genes as input and employed the WikiPathway 2021 gene set (available at https://maayanlab.cloud/Enrichr/#libraries, under the name WikiPathway_2021_Human). The genes were ranked based on the metric (-log10(p-value)/sign(log2FC)). In addition, we set the `fraction` as matched_size/geneset_size, `num` as matched_size, and `log` as -log10(fdr+0.0001). To visualize the pyGSEA results, the ` omicverse.bulk.geneset_plot` function was applied.

In the pyWGCNA analysis, the genes' Median Absolute Deviation (MAD) was calculated using the `statsmodels.robust.mad` method, and the top 5,000 genes with the highest MAD values were selected to construct co-expression modules. Default parameters were used throughout the analysis. The soft threshold for calculating the co-expression network was set to 5, resulting in the identification of a total of 12 modules using the dynamiccuttree approach. We further calculated the Differential Expression Gene (DEG) rate for each module, defined as the number of differentially expressed genes in a module divided by the total number of modules. Module 4 displayed the highest DEG rate, and we identified the gene of interest, APP, within module 5. Visualizations of these two modules were created using the `plot_sub_network` function.

All of the aforementioned analyses were conducted on a computer equipped with an NVIDIA GeForce RTX 2080Ti GPU.

### Supplementary Note 5

#### *OmicVerse provides a multi-type pipeline for single-cell RNA-seq analysis*

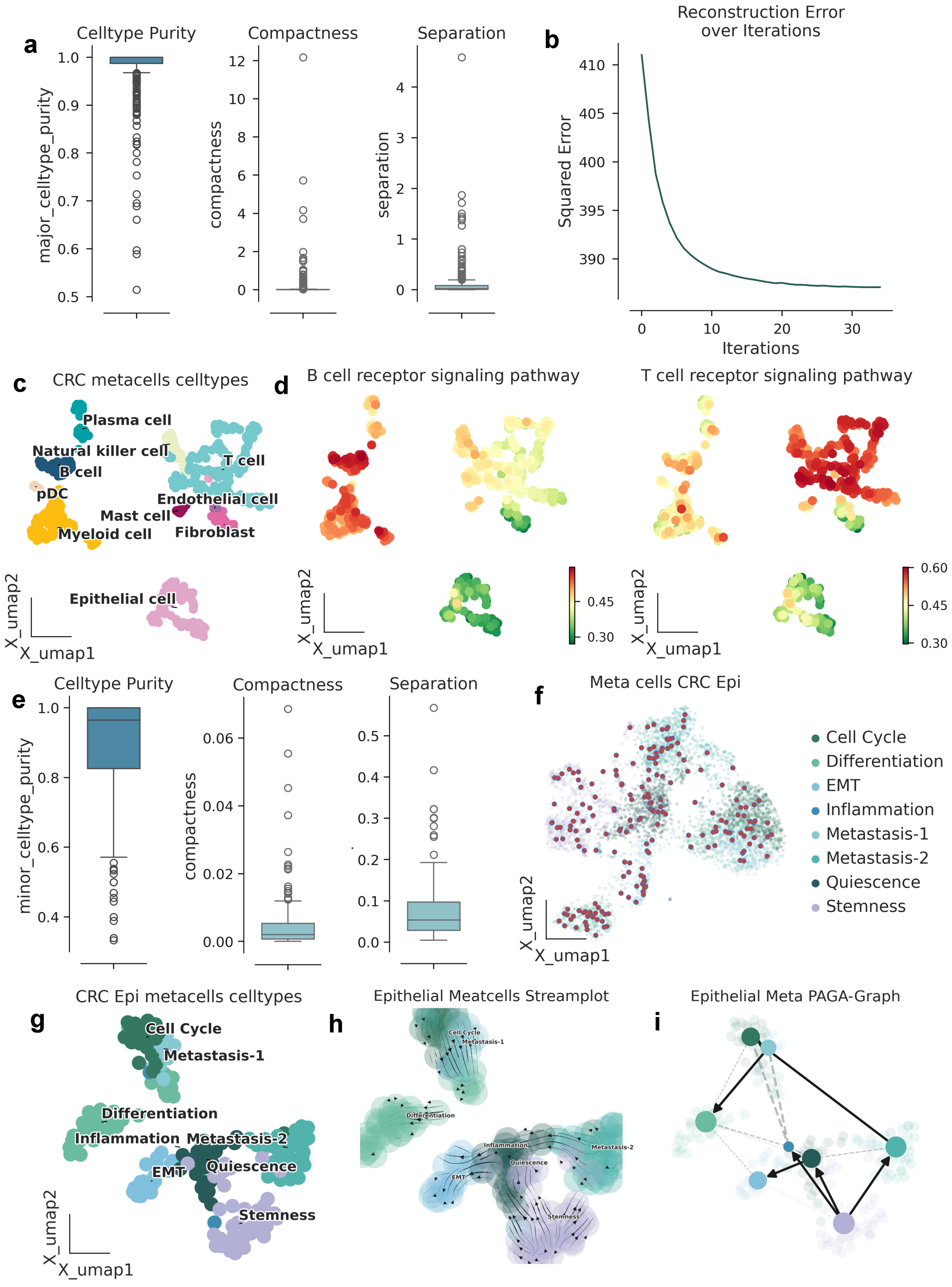

Extended Data Fig 7 | Training Process and Analysis of Metacells

(a) - Distribution of cell type purity, indicating the frequency of the most represented cell type within each metacell. Higher purity signifies a more accurate metacell. The boxes and lines represent the interquartile range (IQR) and median, respectively, while the whiskers extend up to ±1.5 times the IQR. Metacell compactness (average diffusion component standard deviation) and separation (distance between the nearest metacell neighbor in diffusion space; see Methods) are assessed in the CRC single-cell profile.
(b) - Reconstruction error of the CRC data matrix for all cells.
(c) - UMAP plot illustrating single-cell RNA sequencing (scRNA-seq) data of metacells in the CRC, color-coded by cell type annotations.
(d) - UMAP plot representing CRC data and color-coded by pathway enrichment AUCell scores (B cell receptor signaling pathway on the left, T cell receptor signaling pathway on the right).
(e) - Distribution of cell type purity, metacell compactness, and separation in the epithelial CRC profile.
(f) - UMAP visualization highlighting metacells in the raw CRC data.
(g) - UMAP plot illustrating single-cell RNA sequencing data of metacells in the CRC, color-coded by automated annotation of cancer cell subpopulations by pySCSA.
(h) - UMAP plot illustrating the differentiation trajectory of metacells in the CRC profile.
(i) - Cell state transfer directed graph within the trajectory of the PAGA graph in the CRC.

#### *Methods*

The colorectal cancer dataset, corresponding to single-cell RNA sequencing (scRNA-seq) data, was sourced from the Gene Expression Omnibus (Accession ID: GSE178318), as initially reported in a Nature Genetics publication. Associated metadata were likewise collected from the National Center for Biotechnology Information (NCBI) using the same Accession ID. The scRNA-seq data underwent several preprocessing steps. Firstly, low-abundance genes were filtered (min_genes=200), and cells with low expression (min_cells=3) were identified for subsequent filtering. Additionally, a double-cell filtering process was implemented. Following this initial filtering, the data underwent normalization, logarithmic transformation to ensure suitability for downstream analyses.

During the cell annotation process, the 'leiden' algorithm was employed for clustering. For the cell type annotation with pySCSA, the 'celltype' was set as 'cancer,' the 'target' as 'cellmarker,' and the 'tissue' as 'All.' In cases where manual annotation was required, cell markers were sourced from references17, specifically chosen for their relevance to colorectal cancer (CRC). To evaluate the performance of the pySCSA annotation, cells of the same types were merged to match the manual annotation process. The F1 score was computed using 'from sklearn.metrics import f1_score' to assess the annotation quality.

In the pathway (genesets) enrichment analysis, we selected T cell/B cell receptor signaling pathway from the KEGG database. The `omicverse.single.geneset_aucell` was used to calculate the geneset score in all CRC cells.

As for the metacell analysis, we used ` omicverse.single.SEACells` to train the metacell prediction model from the CRC single-cell data file and used `summarize_by_soft_SEACell`, setting the cell type as a parameter. After acquiring the metacells, we also performed pathway enrichment analysis in the same way as before. We also analysed cancer cell subpopulations, we first extracted epithelial cells and cancer stem cells from all cells of the CRC, a total of 11,410 cells. We then used PySCSA again for automatic annotation, except this time our `target` parameter was set to `cancersea`. And in the trajectory analysis module, pyVIA was used to infer the trajectory of cancer cells, the 'adata_key' was set as 'X_pca,' the 'basis' as 'X_umap,' and the start point was designated as 'Stemness.' We also used the same metacellular approach as above for cancer cells, and we used pyVIA for trajectory inference with the same parameters for metacells as well.

In the context of predicting cell interactions, a subset of 14,000 cells was randomly selected from the raw data and used to train a model employing the 'cpdb_statistical_analysis_method’ of CellPhoneDB. The training parameters were configured with 'iterations' set to 1000 and a 'threshold' of 0.1.

All of the aforementioned analyses were conducted on a computer equipped with an NVIDIA GeForce RTX 2080Ti GPU.

### Supplementary Note 6

#### *OmicVerse performed multi-omics factor analysis with MOFA and GLUE*

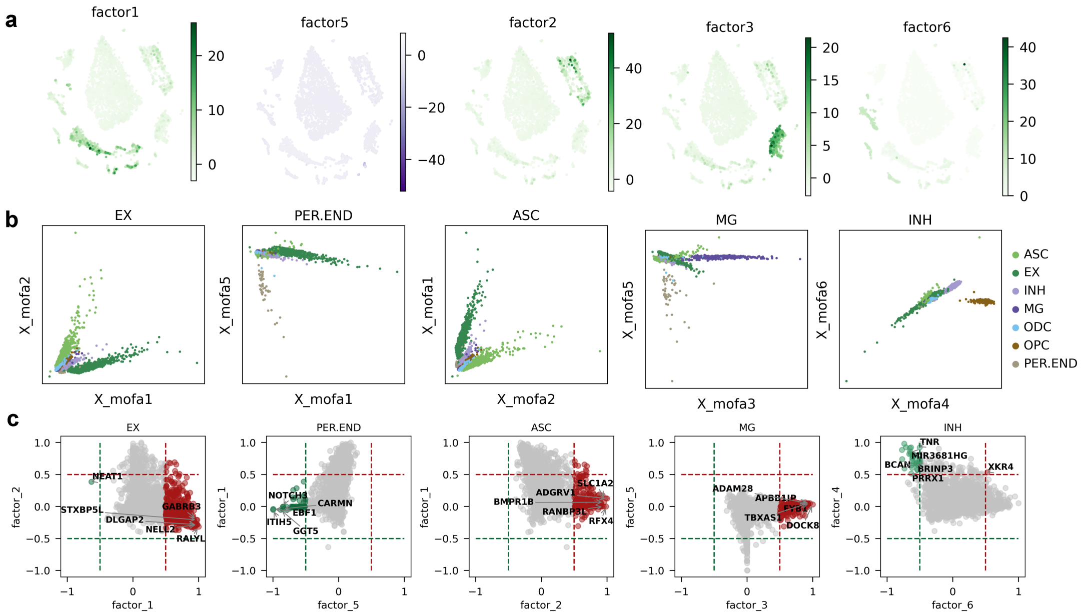

Extended Data Fig 8 | Analysis of Cell Type Variability Using Source Features and Factors

(a) - UMAP embedding illustrating the inferred MOFA factors, where green represents variability derived from positive factor values, and purple represents variability derived from negative factor values.

(b) - Graph displaying the influence of two factors on the source of cell type variability. Each point represents an individual cell, with the horizontal and vertical coordinates indicating the factor values for each cell.

(c) - Co-weight plot of weighted genes for each of the two factors. Each point represents a gene, and the horizontal and vertical coordinates illustrate the weight of a gene on each of the two factors.

#### *Methods*

The data utilized in this study encompassed unpaired single-nucleus RNA sequencing (snRNA-seq) along with single-nucleus Assay for Transposase-Accessible Chromatin using sequencing (snATAC-seq), which was procured from the Gene Expression Omnibus (Accession ID: GSE174367).

For the snRNA-seq data, we undertook a series of preprocessing steps. This entailed normalizing, logging, and scaling the data to ensure uniformity and compatibility for subsequent analysis. We then employed Seurat version 3 (seurat_v3) to identify the top 2000 highly variable genes. Additionally, the top 100 principal component analysis (PCA) embeddings were calculated to capture essential features.

In parallel, the snATAC-seq data underwent a low-expression filter. To facilitate integration, we leveraged the GLUE framework's graph linkage method to transfer the highly variable genes from the snRNA-seq data to the snATAC-seq dataset. The embedding features of cells were derived using latent semantic analysis (LSI).

A critical aspect of our analysis was the construction of a GLUE model, utilizing the original expression matrices from RNA and ATAC data. This model allowed us to capture the relationship between these two data modalities. All preprocessed and analysis step could be found in https://scglue.readthedocs.io/.

Subsequently, GLUE embedding features (`X_glue` stored in .obsm) were calculated for each cell within the snRNA-seq and snATAC-seq histological layers. To connect cells across these layers, we employed `omicverse.single.GLUE_pair` method. Paired cells were used to construct a multi-omics factor analysis (MOFA) model, utilizing `omicverse.single.pyMOFA`. This facilitated the integration and exploration of data across modalities, further enhancing our understanding.

To enhance the visualization of our findings, we conducted a downstream exploration analysis of the MOFA model employing `omicverse.single.pyMOFAART`.

Throughout these analyses, default parameters were utilized. All computational processes were executed on a computer equipped with an NVIDIA GeForce RTX 2080Ti GPU, enabling efficient processing and data integration.

### Supplementary Note 7

#### *Omicverse has a comprehensive and well-established ecosystem in RNA-seq*

The accompanying overview Extended Data figure 9 presents the various stages of the RNA-seq analysis and highlights differences between popular frameworks used for this purpose. omicverse has the same well-established ecosystem for scRNA-seq analysis as seurat, which makes up for the analysis of advanced functions in scanpy, such as automatic annotation of cell types, gene perturbation analysis, cellular interaction analysis, and drug response prediction. In addition, unlike the seurat and scanpy ecosystems, omicverse has a unique Bulk RNA-seq analysis system.

| | | scanpy | Seurat v5 | OMICVERSE |
|---|---|---|---|---|
| Data reading | Start from count matrices | √ | √ | √ |
| Quality control | QC metrices | √ | √ | √ |
| | Doublet removal | √ | √ | √ |
| | Highly Variable Gene Calculating | seurat_v3\|pearsonr | seurat | seurat_v3\|pearsonr |
| Dimensionality reduction | principal component analysis | √ | √ | √ |
| | Latent Dirichlet Allocation (LDA) | ✗ | √ | √ |
| | Visualization | UMAP/TSNE | UMAP/TSNE | UMAP/TSNE/MDE |
| Annotation | Clustering | Leiden/Louvain | Leiden/Louvain | Leiden/Louvain/ GaussianMixture |
| | Find Marker | T test/Wilcoxon | Wilcoxon/logistic regression/ ROC/DESeq2 | T-test/Wilcoxon /DESeq2/COSG |
| | Celltype automatically identity | ✗ | ✗ | pySCSA/MetaTiME/ Celltypist |
| Data integration | Batch correction | Harmony/pyCombat/ FastMNN | CCA/RPCA/ Harmony/FastMNN | Harmony/pyCombat/scanorama/SIMBA/scVI/Mira |
| | Integration with scATAC-seq | ✗ | √ | √ |
| | Metacells/pseudobulk | ✗ | √ | √ |
| | Interpolation from Bulk RNA-seq | ✗ | ✗ | √ |
| | Deconvolution Bulk RNA-seq | ✗ | ✗ | √ |
| Trajectory inference | Diffusion map | √ | √ | √ |
| | Pseudotime Calculated | PAGA graph | monocle3 | pyVIA/Palantir |
| | Gene perturbation analysis | ✗ | ✗ | √(using celloracle) |
| Cell structure | Cell interaction | ✗ | ✗ | √(using CellPhoneDB) |
| | Geneset score | √ | √ | √ |
| | Drug Response predicted | ✗ | ✗ | √(using scDrug) |

Extended Data Fig 9 | Overview of the RNA-seq analysis ecosystem. Python: scanpy and omicverse, R:

Seurat.

#### Data availability

A collection of processed data discussed in this manuscript have been deposited on figshare (https://doi.org/10.6084/m9.figshare.23165636).

#### Code availability

The code to reproduce the experiments of this manuscript is available at https://github.com/Starlitnightly/omicverse-reproducibility. The omicverse package can be found on GitHub at https://github.com/Starlitnightly/omicverse . Documentation and tutorials can be found at https://omicverse.readthedocs.io.

18 Persad, S. *et al.* SEACells infers transcriptional and epigenomic cellular states from si ngle-cell genomics data. *Nature Biotechnology*, 1-12.

19 Hsieh, C.-Y. *et al.* scDrug: From single-cell RNA-seq to drug response prediction.

*Computational and Structural Biotechnology Journal* **21**, 150-157, doi:10.1016/j.csbj.2022.11.055.

20 Gayoso, A. *et al.* scvi-tools: a library for deep probabilistic analysis of single-cell omics data. *bioRxiv* (2021).

21 Ramos, S. I. *et al.* An atlas of late prenatal human neurodevelopment resolved by single-nu cleus transcriptomics. *Nature communications* **13**, 7671, doi:10.1038/s41467-022-34975-2.
